# Shifts in the population-genetic landscape of the ciliate genus Paramecium

**DOI:** 10.64898/2026.05.26.727746

**Authors:** Farhan Ali, Ravina Telkar, Samuel F. Miller, Jiahao Ni, Lydia Bright, John P. DeLong, Kristi L. Montooth, Sascha Krenek, Masahiro Fujishima, Kenji Nanba, Michael Lynch

## Abstract

Ciliates are one of the most ecologically diverse and morphologically intricate unicellular organisms. Despite their evolutionary significance and their prominence in cell-biological research, the population-genetic processes governing their diversification have received remarkably little attention. A fundamental unresolved problem is the existence of geographic isolation among free-living protists and its consequences for species richness. We addressed this issue in the model ciliate *Paramecium* by sequencing genomes of hundreds of isolates collected worldwide, capturing multiple morphological and cryptic species, with multiple populations each. Contrary to previous reports, we found evidence of geographic differentiation in the majority of species. In a few cases, geographic structure became evident when deeply diverging clades in a species were treated separately. This suggests that the biogeographical patterns of *Paramecium* have been shaped by periods of genetic isolation leading to speciation, with rare events of global dispersal realized over its long evolutionary history. Despite being largely isolated, populations were remarkably similar in their effective population size, recombination rate, and efficacy of natural selection. Across species, selection appears to be least effective in *Paramecium aurelia* lineages, and most effective in *P. bursaria*, presumably due to differences in their breeding characteristics. Despite differences among species in the population-genetic environment, patterns of variation across the genome remained consistent. Selective constraints on a core set of genes seemed to have gradually diverged across the species phylogeny. Genes with multiple copies retained from whole-genome duplication events in *P. aurelia* were found to be under relatively relaxed purifying selection. Moving forward, this dataset will serve to test hypotheses on the ecological and cellular complexity of *Paramecium* and beyond.

## Introduction

Darwinian evolution has generated an incredible diversity of biological forms and processes, a substantial share of which is protozoan (Katz 2012; Burki 2014). Studies on *Paramecium*, among a select few, have been key to discovery of fundamental concepts in cell biology such as organelle positioning (Beisson and Jerka-Dziadosz 1999), membrane excitability (Eckert and Brehm 1979), and regulated exocytosis (Plattner 2002). In addition to possessing an elaborate cellular architecture, *Paramecium* are animal-like organisms, that search for food and mates, evade predators, and undergo senescence (Wichterman 1986; Lynn 2010; Van Houten 2023). This makes *Paramecium* an indispensable system in understanding the evolutionary basis of more complex life forms. Despite their evolutionary significance, we have limited understanding of how populations of *Paramecium*, or of any free-living protist, diversify (Lowe et al. 2012; Johri et al. 2017; Weiner and Katz 2021).

Several aspects of ciliate biology make *Paramecium* an exciting and challenging subject for population genetics (Long et al. 2023). Cryptic speciation, where biological species undergo genomic divergence without apparent morphological differentiation, is known in *Paramecium* (Sonneborn 1975) and *Tetrahymena* (Simon et al. 2008) and might be quite common among ciliates (Zhao et al. 2018). This raises issues for studies of population structure where highly diverged populations may belong to different cryptic species (Rynearson and Virginia Armbrust 2004; Koester et al. 2010; Rengefors et al. 2017). Cell multiplication through binary fission is punctuated by occasional but obligatory sexual reproduction, which, in species of *P. aurelia*, can take the form of self-fertilization (autogamy) (Sonneborn 1957). The consequences of these differences in breeding systems for population structure and evolutionary potential, first explored by Sonneborn (1957), can now be examined from a genomic perspective.

*Paramecium* genomes are highly streamlined, gene-rich, and have exceptionally low mutation rates per cell division (Sung et al. 2012; Long et al. 2018). These observations are consistent with large effective population sizes (*N_e_*) for *Paramecium* (Snoke et al. 2006), and have been interpreted in the context of the drift-barrier hypothesis, which predicts that selection can reduce mutation rates and genomic excess more efficiently in species with large *N_e_* (Lynch et al. 2023). Whole genome assemblies for several *Paramecium* species have revealed a history of multiple whole-genome duplications in the divergence of *Paramecium* (Aury et al. 2006; Gout et al. 2023; Ni et al. 2025). The availability of genome sequences for multiple closely related species with varying levels of gene duplication presents an ideal dataset for investigating the evolutionary fate of paralogous genes.

An organism like *Paramecium* that inhabits inland water bodies, does not form cysts, or survive long periods of desiccation, should not be globally distributed (Landis 1988). And yet, common morphological species of *Paramecium* – *P. aurelia*, *P. caudatum*, *P. multimicronucleatum*, *P. bursaria* - are cosmopolitan (Fokin 2010). Several cryptic species of *P. aurelia* also appear to be globally distributed (Przyboś and Tarcz 2018). If dispersal over long distances is sufficiently rare, the degree of genetic differentiation among populations is expected to increase with geographic distance. Previous population genomic work from our group did not support this isolation-by-distance hypothesis (Johri et al. 2017), but strong conclusions could not be drawn due to sampling limitations; only five species – *P.caudatum*, *P. multimicronucleatum*, *P. biaurelia*, *P. tetraurelia*, *P. sexaurelia* – were sampled, with only 10-13 isolates each.

To test whether *Paramecium* populations are geographically differentiated, we sequenced whole genomes of hundreds of *Paramecium* isolates sampled worldwide - covering four morphospecies, with multiple cryptic species and multiple populations for the majority of species. We asked whether these populations vary in the aspects of population-genetic environment - the effective population size, strength of selection, and recombination rate. Finally, having a set of species separated by moderate-to-short evolutionary distance served to examine the pace of functional divergence in protein-coding genes.

## Results

### Geographic structure among *Paramecium* populations

To elucidate the biogeography of *Paramecium*, we have collected and analyzed the genomes of 279 isolates sampled worldwide (Figure 1A). We used mitochondrial genomes of these isolates, along with known sequences, to delineate broad phylogenetic relationships (Figure 1B). Based on a maximum-likelihood phylogeny of mitochondrial amino-acid sequences, we identified nine well-established species in this collection, *viz.*, *P. primaurelia*, *P. biaurelia*, *P. triaurelia*, *P. tetraurelia*, *P. sexaurelia*, *P. quindecaurelia*, *P. caudatum*, *P. multimicronucleatum*, and *P. bursaria*. Additionally, two clades in this phylogeny corresponded to recently described cryptic species of *P. bursaria* and *P. multimicronucleatum* - *P. tribursaria* and *P. fokini* respectively (Melekhin et al. 2022; Greczek-Stachura et al. 2021). *P. bursaria* comprises five syngens (R1–R5) that form monophyletic clades in phylogenies of mitochondrial (mt) (*cox1*) gene sequences (Greczek-Stachura et al. 2012; Greczek-Stachura et al. 2021), ITS, and SSU rDNA sequences (Spanner et al. 2022). Based on similarity to a reference set of *cox1* mtDNA sequences for the five syngens (Greczek-Stachura et al. 2021), we detected three syngens in our dataset - R1 (*P. primabursaria*), R3 (*P. tribursaria*), and R4 (*P. tetrabursaria*). The mitochondrial amino-acid phylogeny failed to resolve R1 and R4 into distinct clades. In general, this mitochondrial phylogeny offered poor resolution for delineating intraspecific relationships; over 60% of internal branch lengths were estimated at 10*^−^*^6^. Therefore, we used neighbor-joining phylogenies based on silent-site differences in the macronuclear genome. These distance-based trees showed that the majority of isolates from a location cluster together, with a few notable exceptions (Figure S1). Isolates of *P. primaurelia* and *P. caudatum* clustered into two deeply diverging clades, referred to as clade A and clade B in both species (Figure 1C). Four isolates of *P. caudatum* from Watson Lake belonged to two different clades, with two isolates in each. The two isolates within each clade are most closely related to each other *i.e.*, exhibiting geographic structure. These clades correspond to the two clades, A and B, of *P. caudatum* identified in Johri et al. (2017). Similarly, for *P. primaurelia*, Catania et al. (2009) reported a highly divergent strain suspected to be a nascent species. Based on these observations, we treated the two major clades in *P. caudatum* and *P. primaurelia* separately for the subsequent analyses.

**Figure 1:**
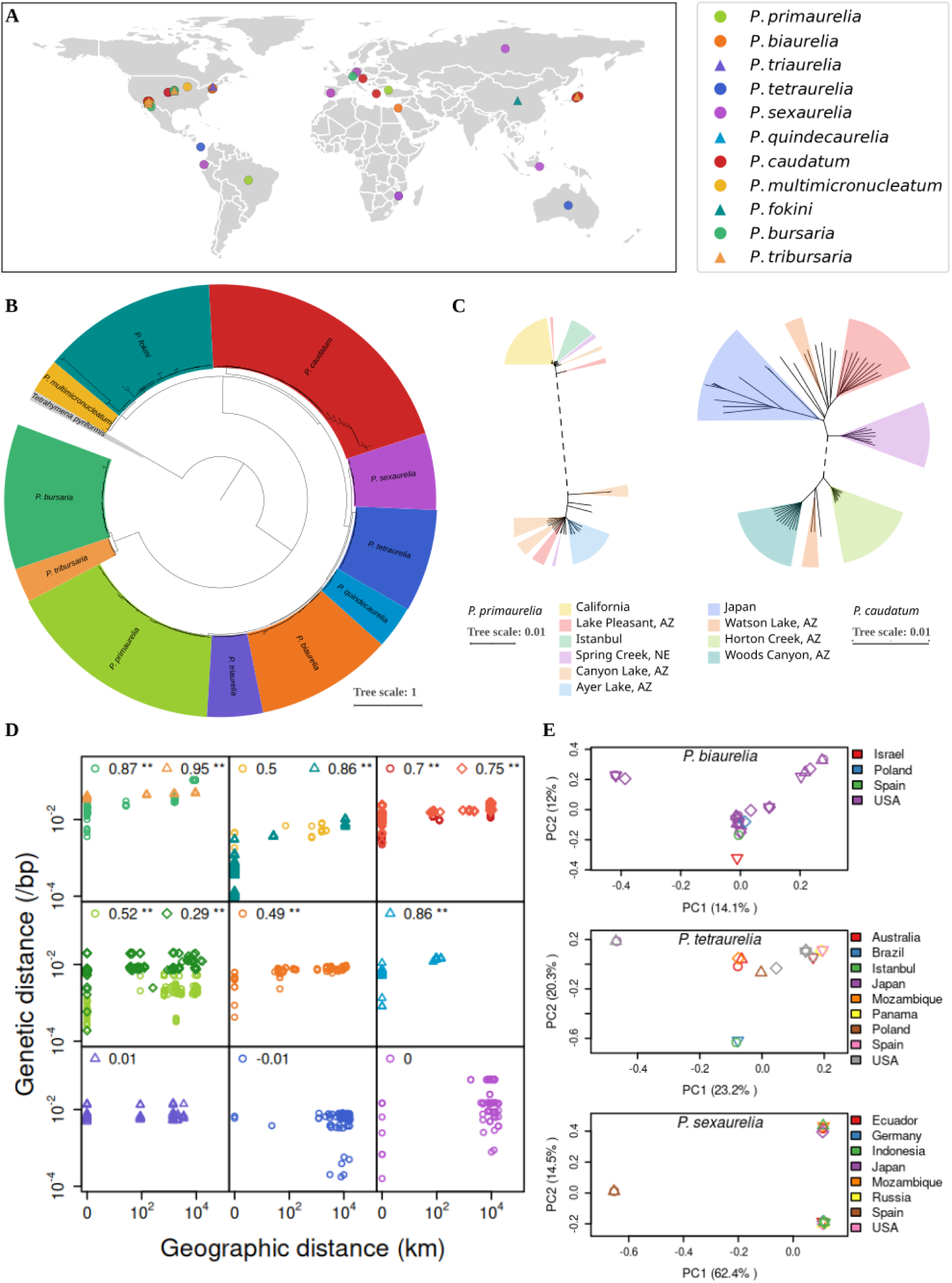
Geographic structure among populations of *Paramecium*. **A)** Sampling locations of *Paramecium* populations marked on an outline of the world map. Populations of different species are shown in colors according to the legend to the right. **B)** Maximum-likelihood mitochondrial phylogeny of concatenated amino-acid sequences of 279 isolates along with reference *Paramecium* mitochondrial genomes, rooted using *Tetrahymena* as an outgroup. **C)** Neighbour-joining phylogenies of *P. primaurelia* and *P. caudautum* isolates based on silent-site distances. Each tree is split into two clades at the central dashed branch. Colored sectors correspond to different populations. **D)** Scatterplots of genetic and geographic distance between individual isolates of a species. Genetic distance between two isolates was calculated as the average of silent-site differences among shared genes. Cryptic species of lineages outside *P. aurelia*, and clades of *P. caudatum* and *P. primaurelia* are shown together in a panel. Species are colored according to the legend in **A**. Spearman’s rank correlation coefficients are mentioned on top of each panel for the respective group of species. ∗∗ indicates *P <* 0.05. **E)** Principal component analysis of sample allele frequencies for isolates of three species - *P. biaurelia*, *P. tetraurelia*, *P. sexaurelia*. Colors correspond to different countries, and isolates from the same location are differentiated using randomly-drawn symbols.

We tested whether the genetic distance between isolates within a species was correlated with the geographic distance between their collection sites, and observed a significant positive correlation for the majority of species, except for *P. triaurelia*, *P. tetraurelia*, and *P. sexaurelia* (Figure 1D). Additionally, for clade A of *P. caudatum* and both clades of *P. primaurelia*, although isolates sampled from the same location showed minimal divergence, genetic distance among isolates from different locations was independent of the geographic distance (Figure S2).

Populations could be geographically isolated even if the genetic divergence between isolates does not scale with spatial distance. We measured genetic differentiation between pairs of populations using *F_ST_*, which is defined as the proportion of variance in sample allele frequencies due to differences among populations in genetic sampling over generations (Weir and Cockerham 1984; Weir and Hill 2002). We estimated sitewise *F_ST_* based on maximum-likelihood estimates of population allele frequencies, computed using GFE (Maruki and Lynch 2015). This approach controls for bias arising from variation in sequencing quality and coverage among samples, and accommodates deviation from the Hardy-Weinberg equilibrium of genotype frequencies.

This analysis was possible for nine pairs of populations from seven species, each with four or more isolates per population. The lowest observed mean *F_ST_* over silent sites was 0.09 for *P. triaurelia* (Table 1). *P. primaurelia* population-pairs had similarly low values - 0.11 and 0.18, whereas values for *P. caudatum*, *P. fokini*, and *P. bursaria*, ranged between 0.27 to 0.59, indicating a moderate to strong population structure. Site-wise *F_ST_* depends on allele frequency as noted previously (Alcala and Rosenberg 2017; Maruki et al. 2022), but the relationship between mean *F_ST_* and minor allele frequency (MAF) varied widely across species (Figure S3). However, in cases where *F_ST_* saturated beyond a certain MAF, the results were consistent with those based on the genome-wide mean. Notwithstanding the evidence for geographic structure among populations, we also observed cases suggesting intercontinental dispersal in *P. biaurelia*, *P. tetraurelia*, and *P. sexaurelia*, based on principal component analysis of individual allele frequencies using PCAngsd (Meisner and Albrechtsen 2018) (Figure 1E). Taken together, these observations present a picture of *Paramecium* biogeography where geographic isolation is the norm, but long-range migration may occur as well in such large populations over a long evolutionary period.

**Table 1:** Pairwise *F_ST_* among populations of seven *Paramecium* species. Populations were defined by sampling location, excluding locations with fewer than four isolates. The mean and standard errors are given for *F_ST_* estimates of 4-fold degenerate sites.

| | Species | Population 1 | Population 2 | Mean $F_{ST}$ | Std. Error |
| --- | --- | --- | --- | --- | --- |
| 1 | *P. bursaria* | Ramsey Canyon Preserve, AZ | Spring Creek Pond, NE | 0.3042 | 0.0057 |
| 2 | *P. caudatum* clade A | Horton Creek, AZ | Woods Canyon, AZ | 0.3600 | 0.0009 |
| 3 | *P. caudatum* clade B | Lake Pleasant, AZ | Fukushima, Japan | 0.3008 | 0.0009 |
| 4 | *P. caudatum* clade B | Lake Pleasant, AZ | Spring Creek Pond, NE | 0.2866 | 0.0006 |
| 5 | *P. caudatum* clade B | Fukushima, Japan | Spring Creek Pond, NE | 0.3047 | 0.0015 |
| 6 | *P. fokini* | Marsh Wren, NE | Spring Creek Pond, NE | 0.5865 | 0.0023 |
| 7 | *P. primaurelia* clade A | CA | Istanbul | 0.1146 | 0.0038 |
| 8 | *P. primaurelia* clade B | Ayer Lake, AZ | Canyon Lake, AZ | 0.1822 | 0.0009 |
| 9 | *P. triaurelia* | NE | Lake Pleasant, AZ | 0.0915 | 0.0006 |

### Variation in the efficacy of selection across the genome and among populations

The degree of genetic polymorphism at distinct classes of genomic sites can reveal differences in selective constraints among these classes and variation in the efficacy of purifying selection across species. Because the majority of mutations are deleterious, the predominant form of selection is expected to be purifying, and more effective selection should lead to lower variation. Given that *Paramecium* genomes are highly streamlined and populations are expected to have large *N_e_*, comparable to estimates in Bacteria (Lynch et al. 2023), polymorphism levels may vary considerably across distinct classes of nucleotide sites. We compared levels of genetic diversity at the following classes of sites: sites of different degeneracies in protein-coding regions, and sites in introns and intergenic regions. We used average pairwise difference, or nucleotide diversity (*π*), as a measure of variation. To control for differences in sequencing quality and coverage among isolates, we estimated site-wise *π* from maximum-likelihood estimates of population allele frequencies (*p*) as *π* = 2*p*(1 − *p*), using GFE (Maruki and Lynch 2015). The reported estimates of species’ nucleotide diversity are site-averaged *π* estimates using all samples of a species.

Across all species, average nucleotide diversity (*π*) is lowest at strictly non-synonymous (0-fold degenerate) sites, and highest at the silent (4-fold degenerate) sites. Overall, *π* increases with *n*-fold degeneracy (Figure S4). Introns and intergenic regions have intermediate levels of diversity (Table 2). This consistency in the ordering of site-classes by polymorphism levels suggests that the relative strength of selection on distinct classes of sites is similar across species. Assuming that selection is sufficiently weak on the 4-fold degenerate (silent) sites such that these evolve neutrally, we used the ratio of nucleotide diversity at non-synonymous and synonymous sites (*π*_0_*/π*_4_) as a measure of the power of selection relative to random genetic drift. Values of *π*_0_*/π*_4_ for all isolates of a species pooled together were lowest in *P. bursaria* and *P. tribursaria*, followed by the species of *P. caudatum* and *P. multimicronucleatum* (Table 2). Species of *P. aurelia* had the highest *π*_0_*/π*_4_, suggesting that the efficacy of purifying selection on non-synonymous sites is at its weakest in *P. aurelia*.

**Table 2:**
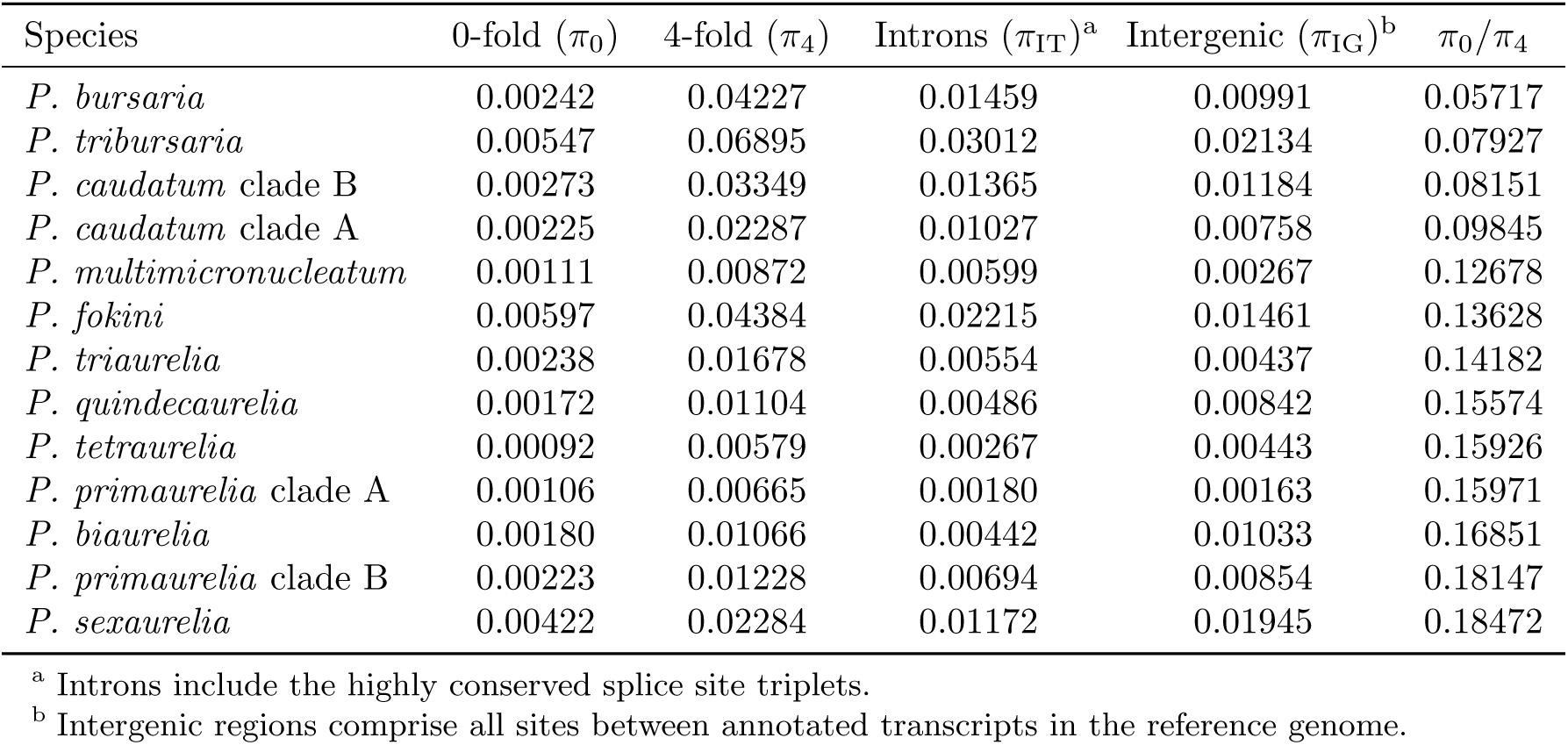
Species-level nucleotide diversity (*π*) at four classes of sites in the *Paramecium* macronuclear genomes. Species are sorted in ascending order of the ratio of non-synonymous to synonymous nucleotide diversity (*π*_0_*/π*_4_).

In the previous section, we observed geographic structure in *Paramecium* populations. This raises the possibility that populations have evolved independently under distinct population genetic environments. In particular, populations at different locations likely differ in their census size, which, in turn, might lead to differences in their local effective population sizes and the efficacy of selection. Upon analyzing each population of a species separately, we observed that *π*_0_*/π*_4_ values tend to be highly similar for different populations within species (Figure 2A).

**Figure 2:**
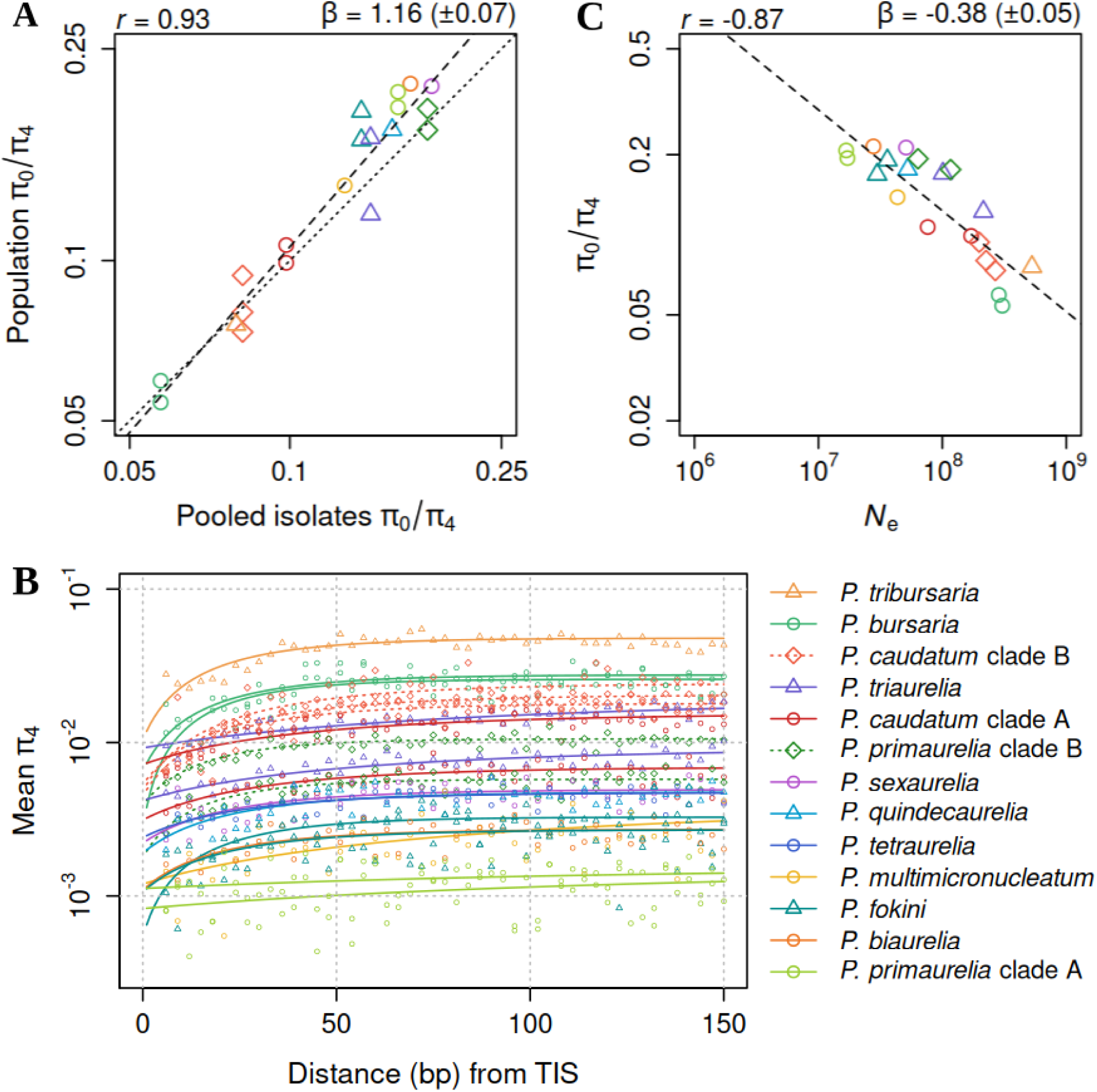
Efficacy of selection across the genome and across species. **A)** The ratio of nucleotide diversity at 0-fold to 4-fold degenerate sites (*π*_0_*/π*_4_) for each population, plotted against species-wide *π*_0_*/π*_4_ estimated using all isolates of the corresponding species. **B)** Average nucleotide diversity across genes at a silent-site position against the distance between the position and the Translation Initiation Site (TIS), excluding intervening introns. Solid lines show asymptotic regression fit for each population, and points show the corresponding raw data. **C)** *π*_0_*/π*_4_ plotted against estimates of *N_e_* derived using the asymptote nucleotide diversity estimated in **B** and the mean of available mutation-rate estimates for three species of *P. aurelia*. Dashed lines in **A** & **C** indicate linear regression fits on a log-log scale. Mean ± standard error (SE) of the regression slopes (*β*), and Spearman correlation coefficients (*r*), are given above the corresponding panels. The dotted line in **A** denotes the one-to-one relationship for species-wide and population-level *π*_0_*/π*_4_ estimates.

In addition to differences across site classes, silent sites showed systematic variation in nucleotide diversity as a function of their relative position. The average nucleotide diversity at silent sites is lowest near the translational start site, asymptotically peaks in the middle, and declines again toward the end. We quantified this relative decline at the start of a gene following Ali (2024). Briefly, we fit a negative-exponential model to estimates of nucleotide diversity at silent-site positions up to a specified distance (*>*= 150 bp) from translation initiation sites (Figure 2B). Details of the model and the determination of the species-specific distance threshold are provided in the Methods. Selection against stable mRNA secondary structures in the translation initiation regions has been proposed to explain their reduced sequence variability observed in Bacteria (Molina and Nimwegen 2008; Ali 2024). Although we do not explore the basis of this constraint in *Paramecium*, the mode and strength of this form of selection are likely conserved across species. Under this assumption, differences among species in the extent of loss in silent-site diversity at the start of genes may be indicative of differences in their effective population size.

Direct estimates of *N_e_*can be obtained based on the expectation of nucleotide diversity at neutrally evolving sites under the infinite-sites model: E(*π*) = 4*N_e_µ*. We used the asymptote of silent-site diversity as the closest approximation of the expected neutral diversity. Mutation rates (per base-pair per generation) have been estimated for three species - *P. biaurelia*, *P. sexaurelia*, and *P. tetraurelia*. Considering the low variability in mutation rates among these species, we approximated mutation rates for other species as the mean of the three experimentally determined estimates, *i.e.*, 2.26 × 10*^−^*^11^. The resulting estimates of *N_e_* ranged from 17 million to 528 million, and negatively correlated with *π*_0_*/π*_4_ (*r* = −0.87*, P <* 2.2 × 10*^−^*^16^) (Figure 2C). This correlation was not a statistical artifact as it persisted even when different populations were used to measure *N_e_* and *π*_0_*/π*_4_ (Figure S5). In conclusion, analysis of nucleotide-diversity patterns across site-classes and species revealed that natural selection is most effective in *P. bursaria*, intermediate in *P. caudatum* and *P. multimicronucleatum*, and least effective among species of *P. aurelia*.

### Recombination as a modulator of the efficacy of selection

Evolutionary fates of mutations are governed by the frequency of recombination. Neutral diversity can be reduced due to physical linkage with selected sites under background selection (Charlesworth et al. 1993; Nordborg et al. 1996), or interference selection (Comeron and Kreitman 2002). Infrequent recombination may lead to linkage disequilibrium (LD), which, in turn, reduces the effectiveness of selection. The degree of linkage in a diploid genome can be measured as the correlation between the heterozygosity of pairs of sites (Lynch 2008). This measure of an individual’s LD - zygosity correlation (Δ) - has the advantage over standard population-level measures of LD (eg, 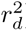) in that it is unaffected by allele frequencies. Δ decays with distance (*d*) between sites in the presence of recombination, as can be seen from the following approximate expression for Δ derived under the infinite-sites model assuming Wright-Fisher population (Equation 8 in Lynch et al. 2014).

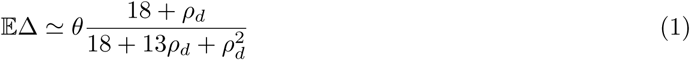

where *ρ_d_* = 4*N_e_cd* is a function of the physical distance (*d*) between sites, and *c* is the recombination rate per base-pair. *θ* = 4*N_e_µ* is the population-scaled per-base mutation rate, which is equal to heterozygosity (*π*) under the infinite-sites model. Because the probability of recombination may be negligible between adjacent sites, 𝔼Δ → *θ* as *d* → 1. On the other hand, 𝔼Δ → *θ/ρ_d_* = *µ/*(*cd*) with *ρ_d_* → ∞.

We used mlRho (Haubold et al. 2010) to estimate Δ between pairs of sites separated by up to 5000 bp for each individual. Overall, Δ declined with the distance between sites in the majority of individuals and species, as expected (Figure 3A). Strikingly, however, Δ values at distances in multiples of 3 (*d*_3_) were highly elevated relative to the rest of the profile (*d*_0_). Because Δ ∝ *θ*, this might reflect higher heterozygosity at third codon positions combined with a high coding density of *Paramecium* macronuclear genomes. Accordingly, Δ profiles for these two sets of distances (*d*_3_*, d*_0_) merge into one with increasing distance as third-codon position pairs from different genes and exons are not necessarily separated by distances in multiples of 3. This may also explain an apparent increase in Δ at higher *d*_0_ distances in *P. bursaria*. Furthermore, the relative increase in Δ at *d*_3_ distances was the highest in *P. bursaria* and, more broadly, was higher among species outside of the *P. aurelia* complex. Although this pattern was present in the Δ decay profiles of *Daphnia pulex* populations too, it was barely noticeable (Lynch et al. 2022). Together, these observations suggest that the striking pattern of wide differences in Δ for different sets of distances in *Paramecium* species reflects their high *N_e_*.

**Figure 3:**
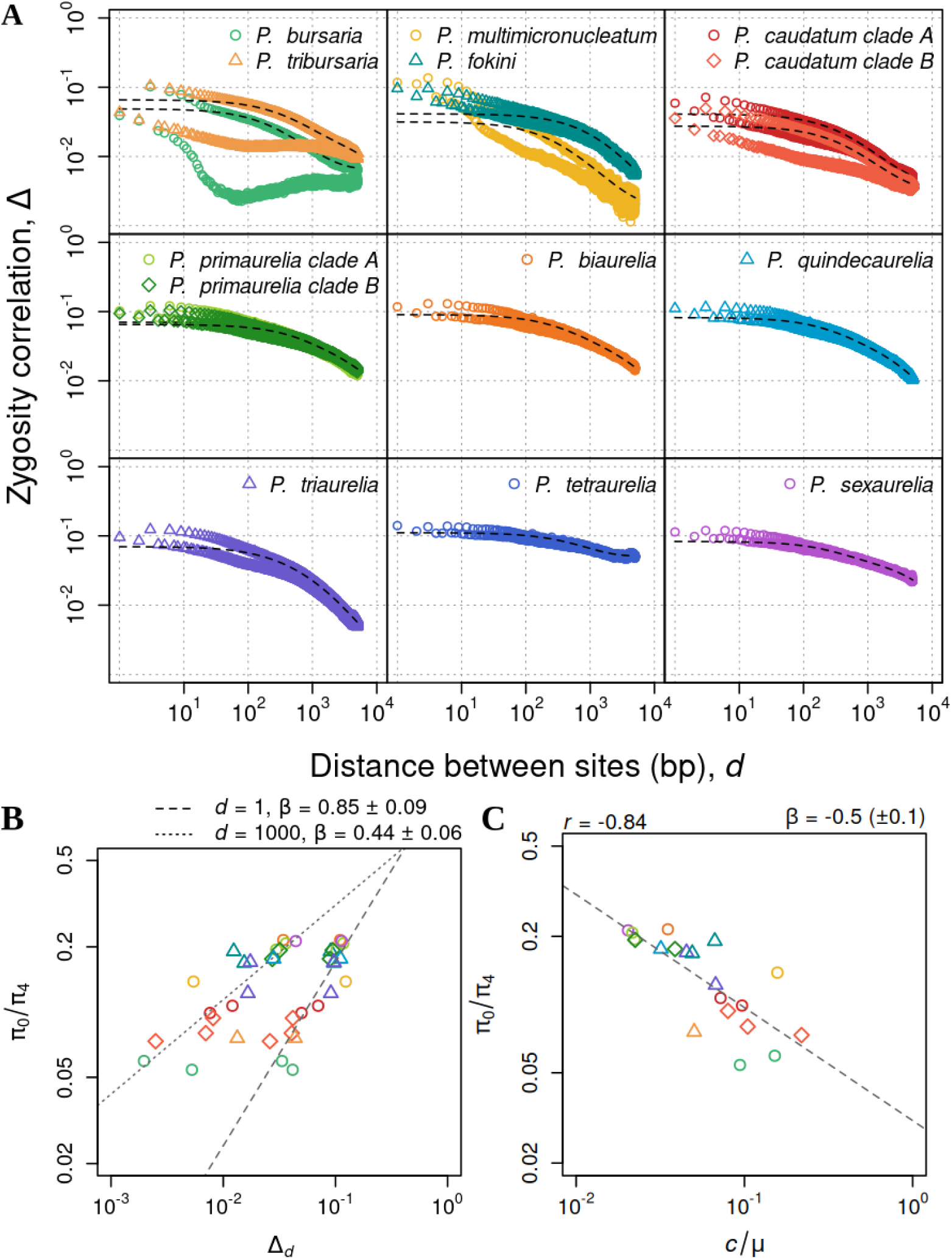
Linkage disequilibrium and the efficacy of selection. **A)** Average zygosity correlation (Δ) across individuals, for pairs of sites separated by physical distance (*d*) in base-pairs, for each species. Closely related species are grouped into separate panels for ease of visualization. Dashed lines show the least-squares fit of Equation 8 from Lynch et al. (2014) to the average Δ decay profile for *d* ∈ (200, 5000), in multiples of 3. **B)** The ratio of nucleotide diversity at 0-fold to 4-fold degenerate sites (*π*_0_*/π*_4_) for each population, plotted against the average Δ across the corresponding individuals at distance *d* ∈ {1, 1000}. Mean ± SE of the linear regression slopes on a log-log scale for different values of *d* are given in the legend. **C)** *π*_0_*/π*_4_ plotted against the ratio of recombination rate to mutation rate (*c/µ*) obtained by dividing the median least-squares estimates of *ρ*_1_ = 4*N_e_c* and *θ* = 4*N_e_µ* from individual Δ decay profiles for each population. Dashed lines show the linear regression fit on a log-log scale, with the corresponding slope (mean ± SE) shown at the top right, and Spearman’s rank correlation (*r* = −0.795*, P* = 3.6 × 10*^−^*^5^) at the top left of the panel.

Δ is a measure of an individual’s linkage disequilibrium (LD). A high LD may result from low *N_e_*, low recombination rate (*c*), or a combination thereof. The efficacy of selection is expected to be lower in species with higher LD. To test this, we used the ratio of population-wide nucleotide diversity at 0-fold to 4-fold degenerate sites (*π*_0_*/π*_4_) as a proxy for the efficacy of selection, with higher values indicating less effective selection. We found *π*_0_*/π*_4_ of a population to be positively correlated with its average Δ across corresponding individuals (Figure 3B), for Δ measured at *d* = 1 (Spearman’s rank correlation, *r* = 0.832*, P* = 2.99 × 10*^−^*^07^, linear-regression slope on a log-log scale, *β* = 0.85 ± 0.09). Because recombination reduces the linkage between sites separated by longer distances, the rate of increase in *π*_0_*/π*_4_ with Δ was lower for the average Δ measured at *d* = 1000 (*β* = 0.44 ± 0.06), although still significant (*r* = 0.898*, P* = 9.65 × 10*^−^*^07^). This confirms that higher LD is associated with lower efficacy of selection across *Paramecium* species, modulated by rates of recombination.

To estimate recombination rates for *Paramecium* species from the patterns of decay in Δ with distance between sites, we used the approach presented in Lynch et al. (2014). This approach accounts for the fact that recombination may occur through non-reciprocal gene conversion in addition to meiotic crossovers by expressing the total recombination rate (*ρ_d_*) between two sites separated by distance (*d*) as

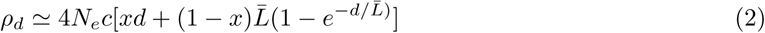

where *x* is the fraction of recombination events accompanying crossovers, and the second term in the bracket is for gene-conversion events with average tract length, *L̅*. By fitting Equation 1 (Equation 8 in Lynch et al. (2014)) to a Δ decay profile using a least squares method, and substituting *ρ_d_* from Equation 2 (Equation 13a in Lynch et al. (2014)), these four recombination parameters (*x, L̅, θ, ρ*_1_) can be estimated. Figure 3A shows the best-fitting curve to the average Δ profile at *d*_3_ distances in each species. *d*_0_ distances were excluded from the curve-fitting because Δ profile at *d*_0_ in several species did not align with the expected curve under the model. Similarly, as seen in Lynch et al. (2014), pairs of sites separated by short distances show higher LD than the expectation under this model and, therefore, were excluded from the analysis using a threshold distance *d* ≥ 200.

We applied this method of estimating recombination parameters to individual Δ profiles in each species. Table 3 reports the median values of parameter estimates across species. In addition to the four parameters, we calculated the relative rate of recombination to mutation (*c/µ*) using estimates of *ρ*_1_ = 4*N_e_c* and *θ* = 4*N_e_µ*. Recombination rate estimates were the highest in *P. multimicronucleatum* (mean *ρ*_1_ = 0.0051 ± 0.0006*, c/µ* = 0.166 ± 0.012), followed by the other outgroup species, and lowest in species of *P. aurelia*, eg, *ρ*_1_ = 0.0016 ± 0.0002*, c/µ* = 0.016 ± 0.003 in *P. tetraurelia*. Thus, recombination rate (relative to mutation rate) varied by an order of magnitude across species. *π*_0_*/π*_4_ was also negatively correlated with *c/µ* (Figure 3C), confirming that recombination-rate variation modulates the efficacy of selection across *Paramecium*.

**Table 3:**
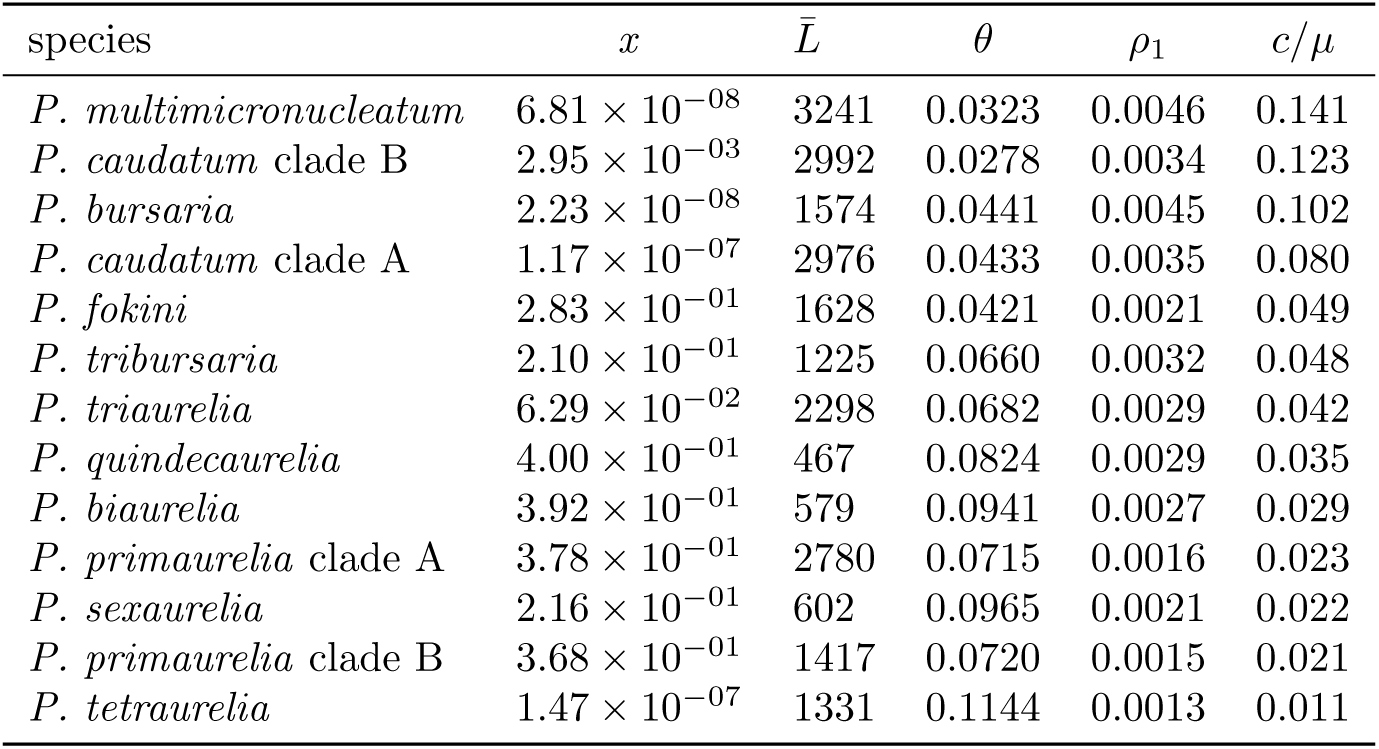
Estimates of recombination parameters from profiles of decay in Δ with physical distance between sites. Reported estimates are median estimates based on the least-squares fit of Equation 8 from Lynch et al. (2014) to individual Δ profiles in each species. Species are sorted in descending order of the ratio of recombination rate to mutation rate (*c/µ*).

### Conservation and divergence of functional constraints across species

So far, we have considered variation in the population-genetic environment across *Paramecium* as an explanation for differences in measures of selection efficacy. An alternative explanation for high *π*_0_*/π*_4_ in *P. aurelia* is the presence of numerous genes, arising from whole-genome duplications, in which the additional copies are under relaxed selection. To test this, we compared the ratio of nucleotide diversity at nonsynonymous and synonymous sites (*π_N_ /π_S_*) for orthologous genes across species. We focused on 5,290 orthologous groups present as a single copy in *P. bursaria* and in one or more copies in four *P. aurelia* species for which both a macronuclear genome assembly and population genomics data were available: *P. primaurelia*, *P. biaurelia*, *P. tetraurelia*, and *P. sexaurelia*. We found an overall increase in *π_N_ /π_S_* with the number of gene copies, as suspected, except that genes with eight copies were at least as conserved as single-copy genes (Figure 4A). However, even for single-copy genes, a paired comparison of *π_N_ /π_S_*between *P. bursaria* and *P. aurelia* supports the conclusion that purifying selection is more effective in *P. bursaria*.

**Figure 4:**
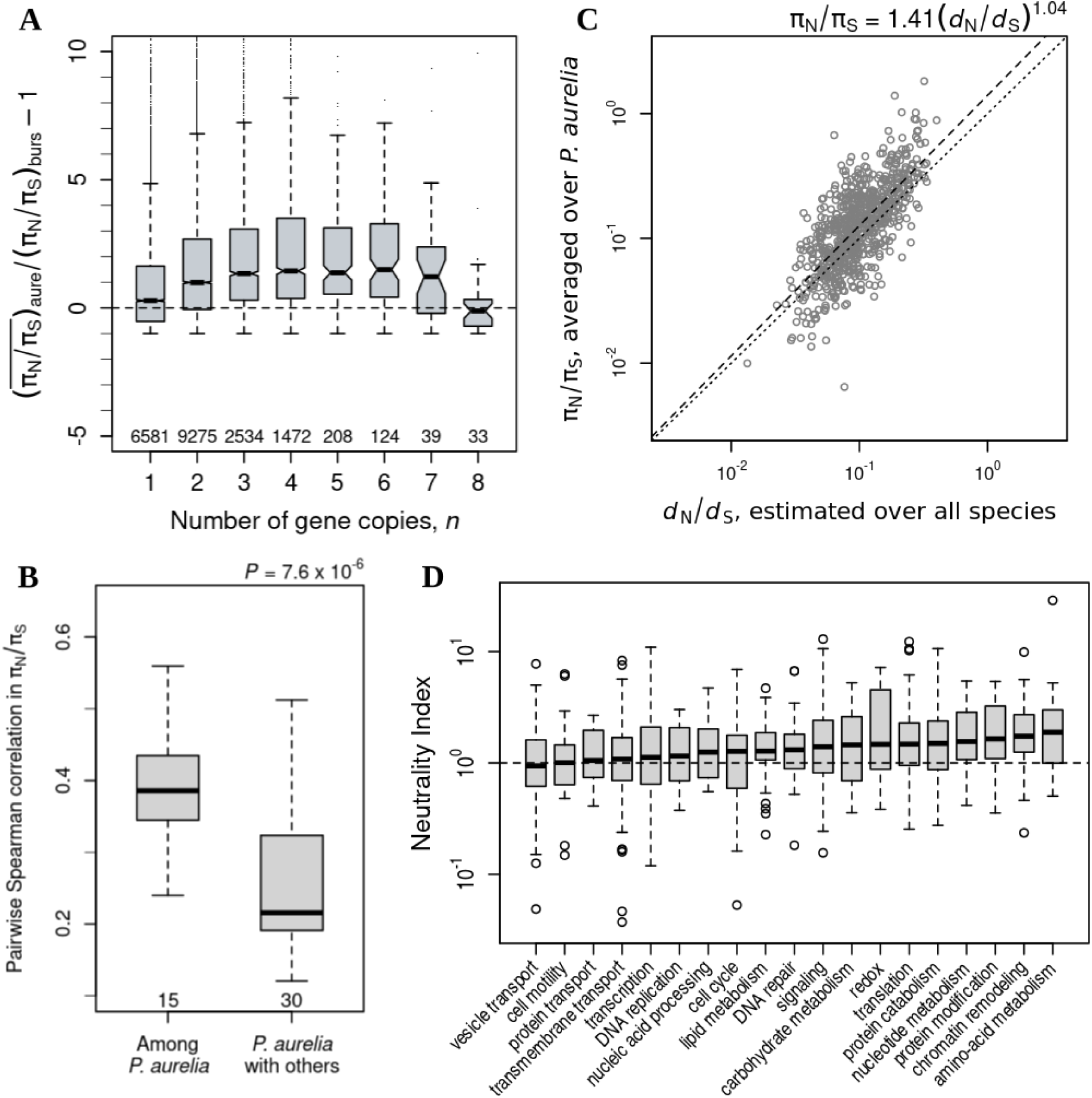
Divergence in selective constraints across proteins and across species. **A)** Relative difference in *π_N_ /π_S_* between *P. aurelia* and *P. bursaria* for single-copy *P. bursaria* proteins, plotted against the number of gene copies (*n*) in a *P. aurelia* species. *π_N_ /π_S_* values for *P. aurelia* are averaged over paralogs within a species. The total number of instances of a gene having *n* number of copies in a *P. aurelia* species is given below each boxplot. The horizontal dashed line at zero marks the expectation of no difference in *π_N_ /π_S_* between *P. aurelia* and *P. bursaria*. **B)** Spearman correlation coefficients for pairwise comparisons of *π_N_ /π_S_* among 6 *P. aurelia* species, and between *P. aurelia* and the 5 outgroup species, for 717 single-copy, core *P. aurelia* proteins. The number of comparisons in each category is given below the respective boxplot. The p-value for a two-sided Wilcoxon rank-sum test is shown at the topright of the panel. **C)** Average *π_N_ /π_S_* across *P. aurelia* species plotted against the overall *d_N_ /d_S_* estimated across the whole phylogeny (including outgroup species) for this set of core proteins. The dashed line shows linear regression fit on a log-log scale, with the best-fit equation (*R*^2^ = 0.496) given above the panel. The dotted line marks the expectation: *π_N_ /π_S_* = *d_N_ /d_S_*. **D)** Distribution of Neutrality Index (NI = (*π_N_ /π_S_*)*/*(*d_N_ /d_S_*)) values of 259 proteins, out of the 717, grouped into 19 functional categories based on Gene Ontology (GO) annotations. The NI for each gene in each species was calculated using a pairwise *d_N_ /d_S_* estimate with respect to its closest orthologous gene (minimum *d_S_*), restricted to cases where *d_S_<* 1. NI values are pooled across species within each category.

Another consequence of *P. aurelia* being gene-rich is that even its single-copy orthologs may be functionally compensated for by excess copies of distantly related genes. Because loss and retention of paralogs proceed independently in different species, the set of single-copy genes experiencing relaxed selection might differ among species. To test this hypothesis, we selected 717 genes that are present in a single copy across all species of the *P. aurelia* complex. Most correlation coefficients for *π_N_ /π_S_* of these genes across species pairs were significantly positive (Figure S6), suggesting that their functional constraints are largely preserved. Nevertheless, *π_N_ /π_S_* values were more strongly correlated among six *P. aurelia* species than between *P. aurelia* and the five outgroup species (Figure 4B), suggesting that selective constraints are less conserved across greater evolutionary divergence.

To test whether orthologous proteins are diverging in function across species as a result of positive selection, we estimated the ratio of non-synonymous to synonymous substitution rates (*ω*) for the above set of genes using a maximum-likelihood approach implemented in PAML (codeml), assuming that the rate is constant across the phylogeny but varies across sites. Note that the mean *ω* for a gene is expected to be less than 1 even under positive selection, because most sites are expected to be selectively constrained. However, if a gene is evolving under positive selection independently in multiple species, then its *π_N_ /π_S_* should be low on average. The average *π_N_ /π_S_* across species for the 717 core proteins was positively correlated with their *ω* values, assessed by linear regression on a log-log scale (*R*^2^ = 0.496*, P <* 2.2 × 10*^−^*^16^) (Figure 4C). Under the fitted model *π_N_ /π_S_* = *κω^α^*, the 95% confidence intervals for the proportionality constant (*κ*) and scaling exponent (*α*) were 1.29–1.55 and 0.96–1.12 respectively. *α* ≃ 1 implies a proportional increase in *π_N_ /π_S_* with increasing *ω*, indicating that high *ω* values predominantly result from relaxed purifying selection. *κ >* 1 suggests that a relatively smaller fraction of non-synonymous polymorphisms reach fixation, presumably due to stronger purifying selection. Taken together, the variation in *π_N_ /π_S_* across these core genes is driven by the strength of selective constraints (Tourasse and Li 2000).

To identify specific instances of positive selection, we turned to a more standard measure of deviation from the neutral model - Neutrality Index (NI = (*π_N_ /π_S_*)*/*(*d_N_ /d_S_*)) (Betancourt et al. 2012). Under (near)neutrality, the relative probability of an allele segregating in the population and going to fixation is the same (NI = 1). Under effective purifying selection, deleterious alleles may segregate at low frequency in the population but do not reach fixation (NI*>* 1). On the other hand, positive selection drives alleles to fixation with limited polymorphism maintained in the population (NI*<* 1). We estimated pairwise *d_N_* and *d_S_* among species for the 717 core proteins using codeml, and calculated NI for each gene in each species with respect to its closest orthologous gene, restricted to cases where *d_S_ <* 1. Again, more than 60% of NI values were positive, suggesting that the purifying selection predominates. Because the individual NI estimates may not be reliable, we sought to detect signs of positive selection over broad functional categories of genes. 354 of these 717 genes had gene ontology (GO) annotations. We classified these genes into 40 categories based on their GO terms description. Of the 19 categories with at least 15 NI values, five had a median NI significantly greater than 1 (*P*_Wilcoxon_ *<* 0.05, multiple-testing corrected), whereas none had a median NI significantly less than 1. Genes associated with transport had among the lowest NI values (Figure 4D). Thus, although some instances of positive selection could be detected, the majority of shared orthologous genes seem to be evolving under selective constraints that are largely preserved across *Paramecium*.

## Discussion

We sequenced and analyzed genomes of hundreds of *Paramecium* isolates collected worldwide, and found that most *Paramecium* populations are geographically differentiated. Although structured by geographic distance, different populations of the same species are similar in their population-genetic parameters. Effective population size, the efficacy of selection, and linkage disequilibrium decay show little variation among populations relative to variation across species. Furthermore, cryptic lineages of the same morphospecies are more similar in these parameters than are lineages of distinct morphospecies. Selective constraints on different site classes and on shared genes are also largely preserved across species, but the effectiveness of selection varies with the strength of recombination and the extent of random genetic drift.

We found the effectiveness of selection to be strongest in *P. bursaria* and weakest among species of *P. aurelia*. Sonneborn (1957) classified species of *Paramecium* based on their breeding characteristics. *P. bursaria* has four mating types, instead of two in *P. aurelia*, has a longer life span (in number of fissions for a clone in the absence of sex) and a relatively long period of sexual immaturity. He argued that these outbreeding features of *P. bursaria* will result in a “common gene pool over a wide range”. This expectation aligns with our observation of high *N_e_* in *P. bursaria* but is contradicted by the strong evidence of population structure. Dini and Nyberg (1993) argued that long periods of asexual growth, such as seen in *P. bursaria*, may lead to local adaptation wherein eventual matings with strangers produce less-fit recombinants, thus reinforcing structure.

Sex in *Paramecium* typically involves physical contact between two cells of different mating types, followed by reciprocal exchange and fusion of haploid micronuclei generated through meiotic recombination. This process is called conjugation. An alternative mode of sexual reproduction, called autogamy, is known in *P. aurelia*, wherein fertilization occurs between two haploid micronuclei of the same individual. Whereas autogamy produces entirely homozygous individuals, conjugation produces a pair of identical heterozygotes. Thus, although population heterozygosity should not systematically differ between the two modes of sexual reproduction, autogamy has the disadvantage of exposing recessive deleterious mutations, which could lead to a decline in *N_e_*if background selection predominates. The relative frequency of autogamy and conjugation in *P. aurelia* populations will depend on mating-type composition. Landis (1988) found only one mating type in *P. tetraurelia* clones isolated from two ponds, and concluded that conjugation is rare in nature for this species. Combined with its ability to undergo autogamy immediately following conjugation, without a period of strictly asexual divisions, *P. tetraurelia* populations may primarily reproduce through autogamy, which might explain the low *N_e_* and limited population subdivision observed in this species.

Despite an apparent lack of geographic subdivision in *P. tetraurelia*, and in *P. sexaurelia* populations, previous studies have reported low F2 survival in crosses between strains of the same species isolated from different geographic regions (Dippell 1954; Stoeck et al. 1998). We found *P. sexaurelia* to have the highest diversity among species of the *P. aurelia* complex, as reported previously (Johri et al. 2017). *P. primaurelia* was the second most diverse species of the group, and did not show isolation-by-distance either. However, we observed two deeply diverging clades in *P. primaurelia* phylogeny. Assuming that these divergent genotypes are reproductively isolated, and analyzing them separately, revealed signs of geographic population structure. Weisse (2008) argued that the ability to detect geographic differentiation will depend on the taxonomic resolution. New cryptic species of *P. aurelia* have been found (Potekhin and Mayén-Estrada 2020) after the original 14 syngens of Sonneborn (1975) were designated as individual species, and there may be several more to be discovered.

The evolutionary history of *Paramecium* is relevant to the problem of geographic differentiation (Preer Jr 1977; De Souza et al. 2020). For example, the synonymous-site divergence of *P. primaurelia* from its closest sister species, *P. pentaurelia*, is 0.069. Assuming mutation rate was as low as 1.22 × 10*^−^*^11^ per site per generation (2 standard deviations below the mean estimate reported for *P. tetraurelia*), and considering 40 generations per year (derived from observed population growth of *P. bursaria* over a 20-day interval (Kosaka 1991), these two species may have diverged as early as 70 million years ago. The partial connectivity within Laurasia during the Late Cretaceous could have facilitated range expansion across suitable freshwater habitats. Since then, populations may have evolved largely independently, leading to speciation, with a combination of continental drift and rare long-distance dispersion obscuring any signal of isolation-by-distance. For species such as *P. sexaurelia* and *P. tetraurelia*, such patterns may become discernible with denser sampling of multiple populations separated by a range of geographic distances.

Notwithstanding strong geographic structure observed in *P. bursaria*, its populations have greater genetic diversity than those of *P. aurelia*. Ecological differences among these species might play a role in determining local *N_e_*. Landis (1988) stated that spatial distribution of *P. aurelia* within a pond is patchy, found almost exclusively at the mud-water interface, whereas *P. bursaria* appears to be widely dispersed owing to its algal endosymbionts and its ability to ingest a variety of prey organisms. Kosaka (1991) tracked population densities of *P. bursaria* in two streams at Hiroshima Prefecture, Japan, and reported median of 0.3 individuals per mL. Hairston (1958), on the other hand, did not report mean densities greater than 0.1/mL for any species of *P. aurelia* sampled from the Huron River, Ann Arbor, over two trips. Assuming a population density of 0.1/ mL spread over a depth of 10 cm, a one-hectare pond will have 10^8^ individuals. Combined with Kosaka (1991)’s observation of two or more equally frequent mating types in *P. bursaria*, its high local *N_e_* may simply reflect a large number of interbreeding individuals.

Recombination is considered to be a major determinant of the efficacy of natural selection in microbes (Price and Arkin 2015), and we have shown this to be true in *Paramecium* as well. Rengefors et al. (2017), in their review of genetic diversity in eukaryotic phytoplankton, argued that prolonged asexual phases will lead to a high LD because the entire genome will act as a single linkage group. Additionally, populations with greater rates of inbreeding show higher LD, eg, selfing populations of *Arabidopsis lyrata* were found to have increased correlation of zygosity (Lucek and Willi 2021). Consistent with these observations, we found recombination rate estimates (*c/µ*), based on Δ) decay profiles, to be lower in *Paramecium* than those reported for obligately sexual vertebrates in Lynch et al. (2014). *c/µ* for *Daphnia pulex*, a facultatively sexual invertebrate, was found to be 0.034, which is intermediate to the range of values seen across *Paramecium* species. Among *Paramecium*, *P. aurelia* may have lower rates of recombination due to reproduction through autogamy, an extreme form of selfing.

Although demographic and genetic factors influence the evolutionary dynamics of different species of *Paramecium*, functional constraints on shared sets of proteins are largely preserved across the range of evolutionary divergence examined in this study. These shared constraints presumably reflect the shared aspects of their biology (Wichterman 1986; Long et al. 2023). Especially among cryptic species, where morphological differences seem non-existent, a pertinent question is whether these species are ecologically equivalent (Hairston 1967; Nyberg 1974; Fišer et al. 2018; S^̌^kaloud and Rindi 2013). Simon et al. (2008), in their review on morphospecies of *Tetrahymena pyriformis*, argued that the commonality of ecological niche is highly unlikely in the face of many cases of molecular differences among cryptic species. Because cryptic species of *Paramecium aurelia* likely emerged over 100 million years ago following a whole-genome duplication, the potential for metabolic divergence despite apparent morphological similarity is significant (McGrath et al. 2014; Gout et al. 2023). Indeed, Sasaki et al. (2006) connected differences in thermal tolerance of syngens of *P. aurelia* to differences in their fatty-acid composition. Although the conservation of selective constraints on shared genes was noted previously (Johri et al. 2017), the inclusion of *P. bursaria* in the present study, and of multiple cryptic lineages outside of *P. aurelia*, revealed the extent and generality of this phenomenon. Further developments on the population and evolutionary genomic dataset for this model ciliate will be invaluable to understanding how variation in gene content translates into cell-biological and, consequently, ecological differences among species.

## Methods and Materials

### Sample collection and culturing

Samples were collected from freshwater habitats with stagnant water and a loamy substrate. Sampling was performed by disrupting the substrate with a bottle/golf ball retriever with a plastic bag attached to the end and scooping up the resulting cloud of water. For larger water bodies, samples were collected at sites separated by 50 m along the shoreline, or from the same site during multiple visits interspersed over months. The sampled water was split into batches within a day of collection by shaking the container and pouring 25 mL into individual Petri dishes, supplemented with 1-3 grains of unsprouted rice.

Isolation was performed within a couple of days after collection, depending on population density. Approximately 5 *Paramecium* were picked out of a Petri Dish and, using a Pasturer pipette with a flame-narrowed stem, were moved into the first well of a 3-well plate filled with either Dryl’s, inoculated wheatgrass medium, or lettuce juice medium. Samples were washed by a single transfer of 3 *Paramecium* through the second and third wells before transferring each isolate into individual wells of a 3-well plate with 200*µ*L of inoculated culture fluid. If clearly distinguishable species were seen within a single Petri dish, *i.*e. *P. tetraurelia* and *P. caudatum*, isolates were selected to transfer both. When photosynthetic *Paramecium* were found in culture, the isolation process was the same except the isolated cells were moved to culture tubes containing 1mL of culture fluid and incubated in light. Cultures were fed every 2-3 days with bacterium *Klebsiella aerogenes* (ATCC 35028) in the above-mentioned media for a week until one surviving isolate (or more if morphologically distinguishable) population is selected for long-term culturing and genome sequencing.

Of the 279 isolates analyzed in this study, 81 were obtained from *Paramecium* collections maintained in Germany (Institute of Hydrobiology – TU Dresden), and in Japan (Yamaguchi University), and three were collected in China (Institute of Evolution and Marine Biodiversity, Ocean University of China). A total of 79 isolates were sampled from Nebraska, New York, Wisconsin, and California. The remaining 116 samples were collected from eight locations in Arizona. Details of the sample locations are provided in Table S1.

### DNA isolation and sequencing

DNA purification was either performed using the MasterPure Complete DNA isolation kit MC89010 (Biosearch) or the MagMax Ultra (Life Sciences A36570) 2.0 kit, run on a KingFisher Flex with a 200*µ*L extraction program setting. Libraries were generated using the Ultra II FS DNA Library kit for Illumina (NEB Ultra II FS DNA Library Prep kit for Illumina E7805 with Multiplex oligos dual index primers sets – E7600, E7780, all New England Biolabs). with a miniaturized protocol (Li et al 2019) with the number of amplification cycles reduced to 6-7 to minimize overamplification problems. These samples were inspected for size on an Agilent Tapestation 4200 using High Sensitivity D-1000 screentapes, and quantified using a Qubit 4.0. Acceptable samples had band peaks between 300-700 bp and at least 500 pg/*µ*L. Pooling was performed with normalization based on peak size and quantity, concentrated and cleaned to produce samples of sufficient concentration, and the pooled libraries were sent for 150bp paired-end whole genome sequencing.

### Species identification

Sequencing reads of all samples were mapped to reference macronuclear genomes of 17 *Paramecium* species available on ParameciumDB (Arnaiz et al. 2019) using BWA-mem (v0.7.17) (Li 2013). The reads unmapped to the nuclear genome with the highest mapping percentage were extracted using SAMtools (v1.17) (Li et al. 2009) and mapped to the genomes of the feed bacterium and of known bacterial endosymbionts. The remaining reads were de novo assembled using SPAdes meta (v3.15.5) (Nurk et al. 2017), and mitochondrial contigs were identified by BLAST search (Altschul et al. 1997) against a custom *Paramecium* cox1 database (Long et al. 2023). The mitochondrial genomes were annotated using the genetic code recommended by Arnaiz et al. (2019) and by BLAST searches against mitochondrial protein sequences from ParameciumDB. Amino-acid sequences were concatenated, aligned using MAFFT (Katoh and Standley 2013), and filtered using BMGE (Criscuolo and Gribaldo 2010). A maximum-likelihood phylogeny was constructed on this multiple-sequence alignment using RaxML assuming the LG+G substitution model (Stamatakis 2014). The final species designation for the new samples was based on their colocalization with the reference samples of known species in the phylogeny. Details of the mitochondrial annotation and phylogenetic reconstruction will be published separately.

### Variant calling and filtering

Variant calling was based on the reads mapped to the reference nuclear genome corresponding to the final species assignments as above. For four species without a reference assembly, the closest available reference was used. BCFtools (v1.16) mpileup was used, along with a multi-allelic (rather than consensus) caller, to identify variants assuming diploidy (Danecek et al. 2021). Maximum read depth was adjusted for a batch of samples to be slightly greater than twice the highest mean depth of coverage. Reads with average mapping quality and base quality below 20 were excluded, and base-alignment quality scores were recalibrated to improve SNP detection around Indels. SNPs within three bp of indels, and Indels separated by less than nine bp were removed. The distributions of variant quality stats - Quality scores, Depth, Mapping quality bias, Base quality bias, Read position bias, Strand bias - of the remaining SNPs were manually examined to identify appropriate thresholds for filtering. Selected SNPs were annotated using locally built SNPEff databases for the corresponding reference genomes (Cingolani et al. 2012). Presence of genes in a sample was evaluated using SAMtools bedcov. Reads with secondary and supplementary mappings, duplicate reads, and reads failing quality checks were excluded. Genes with less than 60% of their length covered by at least one read (*i.e.*, breadth of coverage), or genes with depth of coverage three standard deviations below the mean depth, were excluded. Only the SNPs falling within the selected genes in a sample were retained.

### Estimation of nucleotide diversity

The nucleotide class and degeneracy of a site were determined based on the reference genomes. Non-synonymous and synonymous SNPs were extracted from the SNPEff annotations. Non-synonymous and synonymous nucleotide diversity of a gene in a species were calculated as the average number of non-synonymous and synonymous differences across all pairwise comparisons, respectively, divided by the corresponding reference site counts adjusted for the gene’s mean breadth of coverage across samples. The program GFE was used to estimate population allele frequencies (*p*) based on a maximum-likelihood approach (Maruki and Lynch 2015). Site-wise heterozygosity is estimated as *π* = 2*p*(1 − *p*). This method co-estimates an error rate (*ε*)for each site, which was used to filter sites such that *ε <* 0.05. All variable and invariant sites within each class that met the following criteria were used in the analysis. These were sites detected in at least 70% of individuals, using uniquely mapped reads with mapping quality *>* 25 and base quality *>* 30. Estimates of nucleotide diversity for site classes are averages of *π* over class-specific subsets of sites.

### Population structure analysis

The evidence for geographical population structure in *Paramecium* was analyzed in three ways. First, the correlation between genetic and geographic distance was assessed. Genetic distance between a pair of individuals within a species was defined as the average number of silent-site differences across genes. Geographic distance was computed as pairwise geodesic lengths (in kilometers) using an Earth ellipsoid model (WGS84) in Python-based module GeoPy. Second, the PCAngsd program (Meisner and Albrechtsen 2018) was used to perform principal component analysis (PCA) on a covariance matrix for genotypes at variable sites. Because genotypes are not known with certainty in NGS data, the program uses genotype likelihoods. Silent sites with high-confidence SNPs (*P <* 10*^−^*^6^) that satisfied all the aforementioned criteria were used in the analysis. The third approach was to estimate pairwise *F_ST_* for populations within a species using PSA (Maruki and Lynch 2015) based on the statistical framework of (Weir and Cockerham 1984). PSA reports site-wise *F_ST_* estimates. Sites were filtered using the same thresholds as in the section for estimation of nucleotide diversity. Summary statistics were calculated on the *F_ST_* distribution of the remaining sites.

### Analysis of patterns of silent-site diversity over the gene length

All silent-site positions at a certain distance from the translation initiation site (TIS), up to 500 bp away, were pooled across genes. Mean of site-wise nucleotide diversity was calculated for these subsets of silent sites. An asymptotic regression fit of the following form was applied to the pattern of mean nucleotide diversity over distance (Ali 2024).

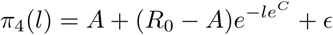

where *A* stands for the asymptote level of diversity and *R*_0_ for the minimum estimate of diversity at position index 0 (TIS). *C* determines the rate of increase in diversity with distance. The error in the fit is additive and independent, with variance inversely proportional to the number of genes for the position. The maximum position used in model fitting for a species was determined through exploratory analysis. A range of distance thresholds was tested and evaluated using two measures: effect size (*L*_eff_= log_2_(*A/R*_0_)) and effect length (*S*_eff_ = ln2*/e^C^*). The profiles of effect size and effect length over distance thresholds were visually examined, and the maximum distance at which these measures appeared to stabilize was selected as the final threshold. Fitted parameters and distance thresholds for each species are provided in the Table S2.

### Linkage disequilibrium analysis

Recombination and linkage disequilibrium analyses were performed using mlRho (v2.9), which estimates correlations of zygosity (disequilibrium at the individual level) while accounting for sequencing errors in a maximum-likelihood framework. Details on the conceptual basis and implementation can be found in Lynch (2008) and Haubold et al. (2010). Briefly, under a neutral Wright-Fisher model, the expected nucleotide diversity at a site is 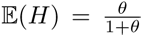 where *θ* = 2*Nµ* is the population-scaled mutation rate. Considering pairs of sites separated by a distance under physical linkage, we have the following expressions for the expected proportion of pairs of heterozygous (H2) sites.

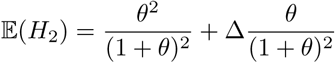

which gives

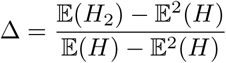

Defining *h_i_* as the zygosity of a diploid individual at site *i*, with *h_i_*= 0 if the site is homozygous and *h_i_* = 1 otherwise, it can shown that

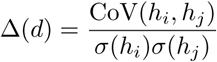

is the correlation of zygosity of two sites *i* and *j* separated by physical distance *d*, such that Δ = 0 gives the expectations for independent sites and Δ = 1 gives the expectations under complete linkage. For any intermediate level of recombination, given by population-scaled recombination rate (*ρ_d_* = 4*N_e_cd*), Δ decreases with increasing *ρ*.

mlRho requires a table (profile) of counts for all four bases at each selected genomic position as input. This profile was generated by selecting high-quality, primary mappings with a minimum mapping quality of 25, a minimum base quality of 30, and positions with at least 5 mapped reads. In averaging Δ across individuals, only valid Δ estimates were used - 0 *<* Δ *<* 1 with limits Δ_min_ *>* −1 and Δ_max_ *<* 1. Recombination parameters were estimated by fitting Equations 1 and 2 (Equations 8 and 13a in Lynch et al. (2014)) to individual profiles of decay in Δ with distance between sites, using a least squares method. The objective function was the mean squared difference in the predicted and observed Δ*_d_* in log space for 200 ≤ *d* ≤ 5000. The fit was optimised using the Nelder-Mead algorithm along with the initial parameter estimates - *θ* = Δ̅, *ρ*_1_ = 10*^−^*^3^*, x* = 0.1*, L̅* = 1000.

### Selection analysis on divergence data

Orthologs of nuclear-encoded proteins across 16 *Paramecium* species were identified using OMA (v2.6.0) (Altenhoff et al. 2019). 39383 orthologous groups were identified at the root level. For polymorphism analysis across paralogs, 5290 gene families with one or more copies in all *P. aurelia* species and a single copy in *P. bursaria* were selected. 717 single-copy core orthologous families in 13 species of *P. aurelia*, that may or may not have an ortholog in the three outgroup species (*P. caudatum*, *P. multimicronucleatum*, *P. bursaria*), and that were also reported in (Gout et al. 2023), were selected for divergence and functional analysis. Clustal Omega (v1.2.4) (Sievers et al. 2011) was used to generate alignments of amino-acid sequences, which were converted to codon alignments using pal2nal (v14)(Suyama et al. 2006) under NCBI codon table no. 6. The codeml program in the PAML package (v4.10.9) was used to estimate the ratio of non-synonymous to synonymous substitutions (*ω*) for each gene under maximum-likelihood models of sequence evolution (Yang 1997). The M8 model was applied to the alignments, which assumes a beta distribution of *ω*(*<* 1) across sites. (Yang et al. 2000). The *ω* estimates for the genes presented in the results section are expectations computed from the beta-distribution parameters (*p, q*), *i.e.*, 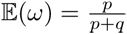. Neutrality Index (NI = (*π_N_ /π_S_*)*/*(*d_N_ /d_S_*)) of a gene in a species was calculated using the pairwise *d_N_ /d_S_* estimate (using codeml) with respect to the closest ortholog (shortest *d_S_*). NI values were only calculated for comparisons with 0 *< d_S_ <* 1; *d_N_ >* 0; *π_N_ >* 0; *π_S_ >* 0.

Functional classification of the 717 genes was performed using Gene Ontology (GO) annotations from the reference *P. tetraurelia* assembly (ASM16542v1), accessed *via* Ensembl Protists. 354 of the 717 genes, with one or more GO terms, were assigned to 40 categories by curating their GO terms’ descriptions (Table S3).

## Acknowledgments

This work was supported by the National Science Foundation (DEB-1927159, DBI-2119963), National Institutes of Health, R35-GM122566-08, and Grant 735927 from the Moore-Simons Project, awarded to Michael Lynch, and by the National Science Foundation grant (IOS #1838098) awarded to John P. DeLong and Kristi L. Montooth.

## Supplementary Material

**Table S2:**
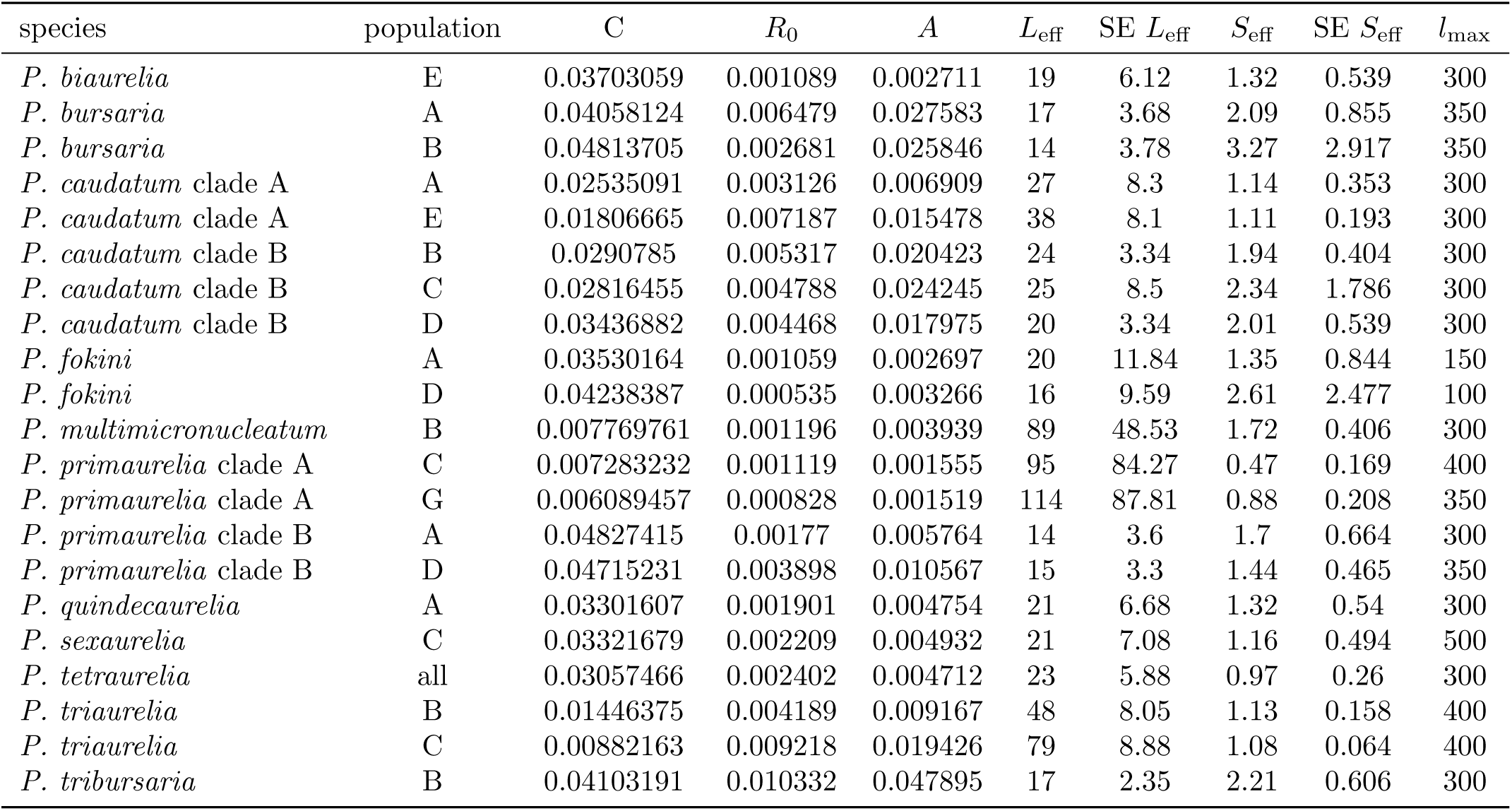
Estimated parameters from asymptotic regression of change in mean nucleotide diversity at silent sites with distance from translation initiation site, excluding introns. Parameters are as defined in the Methods. *l*_max_ is the maximum distance used for the best-fitting model.

**Figure S1:**
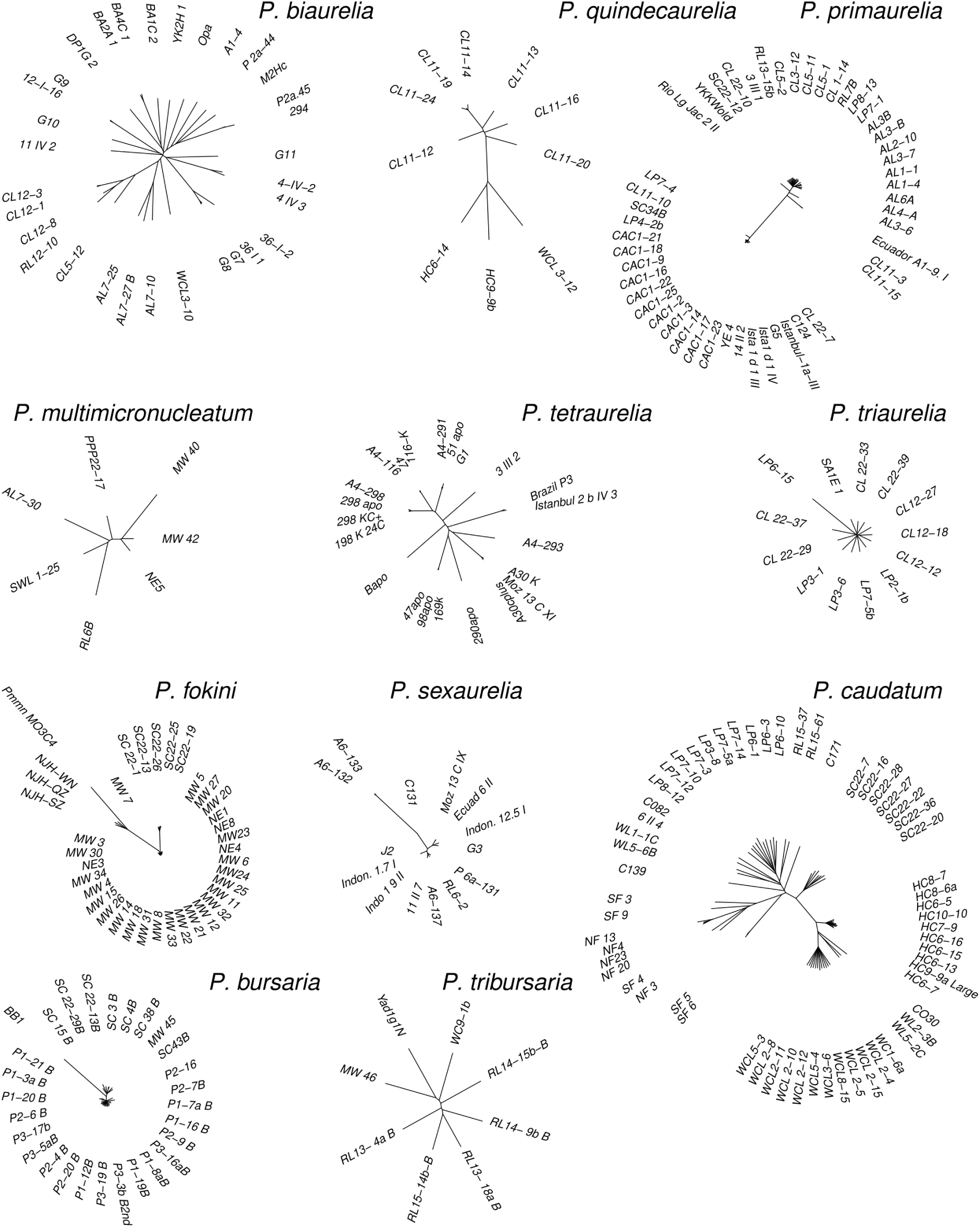
BioNJ distance phylogenies of individual species based on silent-site differences in the macronuclear genome.

**Figure S2:**
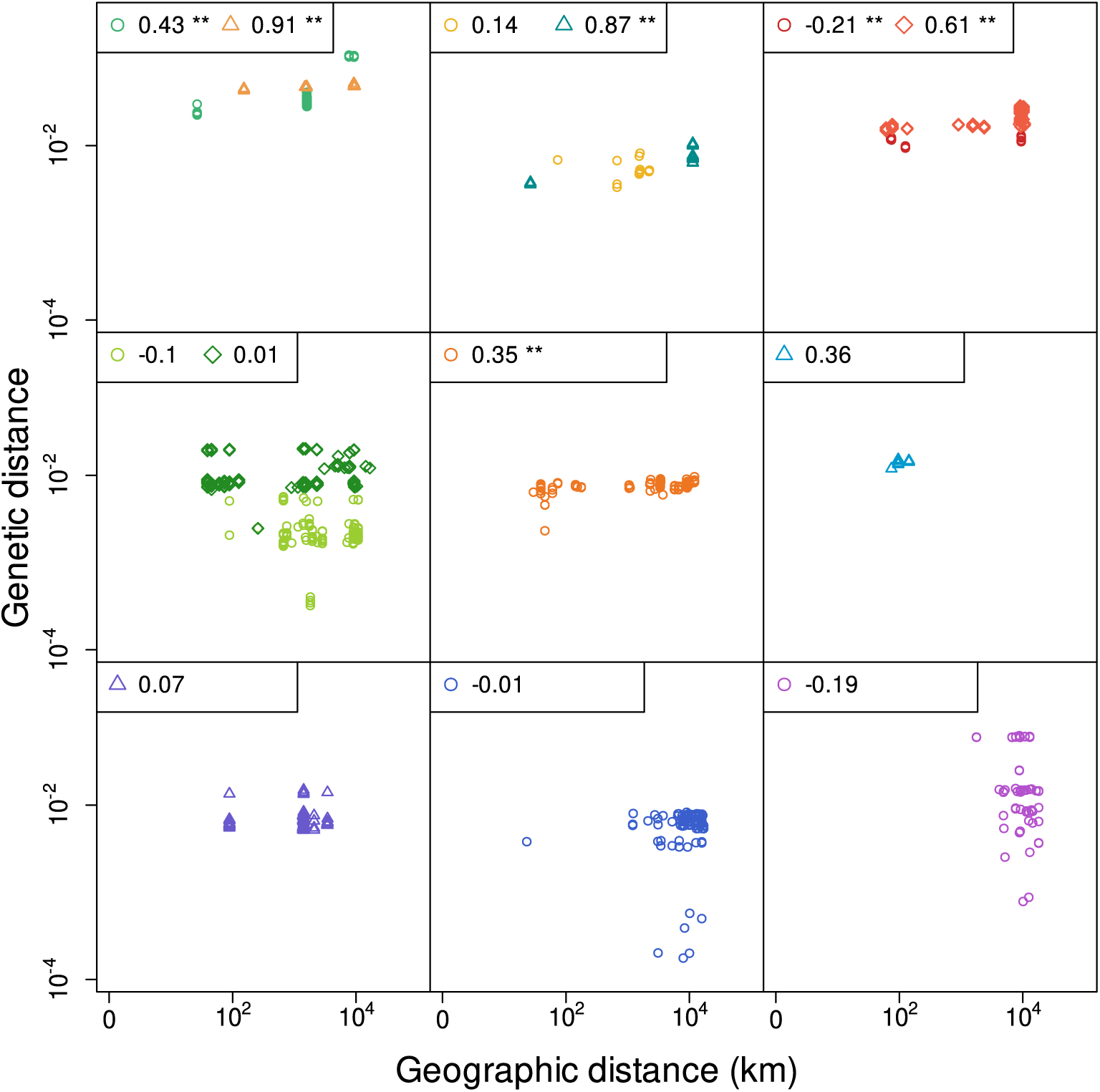
Pairwise genetic distance, based on silent sites differences in the macronuclear genome, among samples isolated from different locations against geographic distance between the sites of collection. Species are shown in different panels and colors, as in the main Figure 1D. Shown on the top-right of each panel are Spearman correlation coefficients, with asterisks for *P <* 0.05.

**Figure S3:**
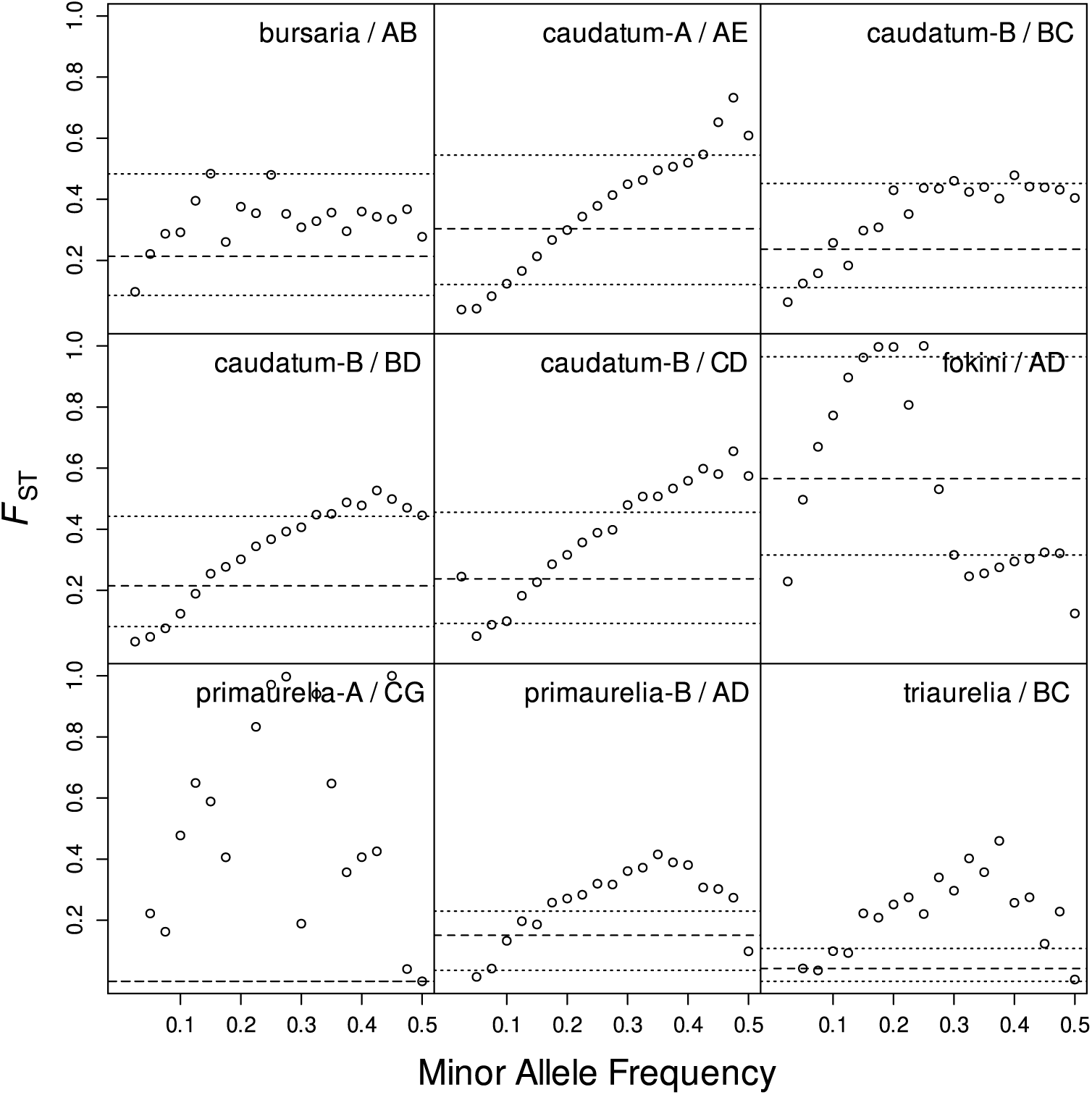
Change in GFE estimates of site-wise *F*_ST_ with minor allele frequency (MAF) for pairs of populations in different species. Dashed lines show median values and dotted lines show the 25th and the 75th percentiles *F*_ST_ over all sites. Points show the mean *F*_ST_ over non-overlapping bins of MAF of size 0.025.

**Figure S4:**
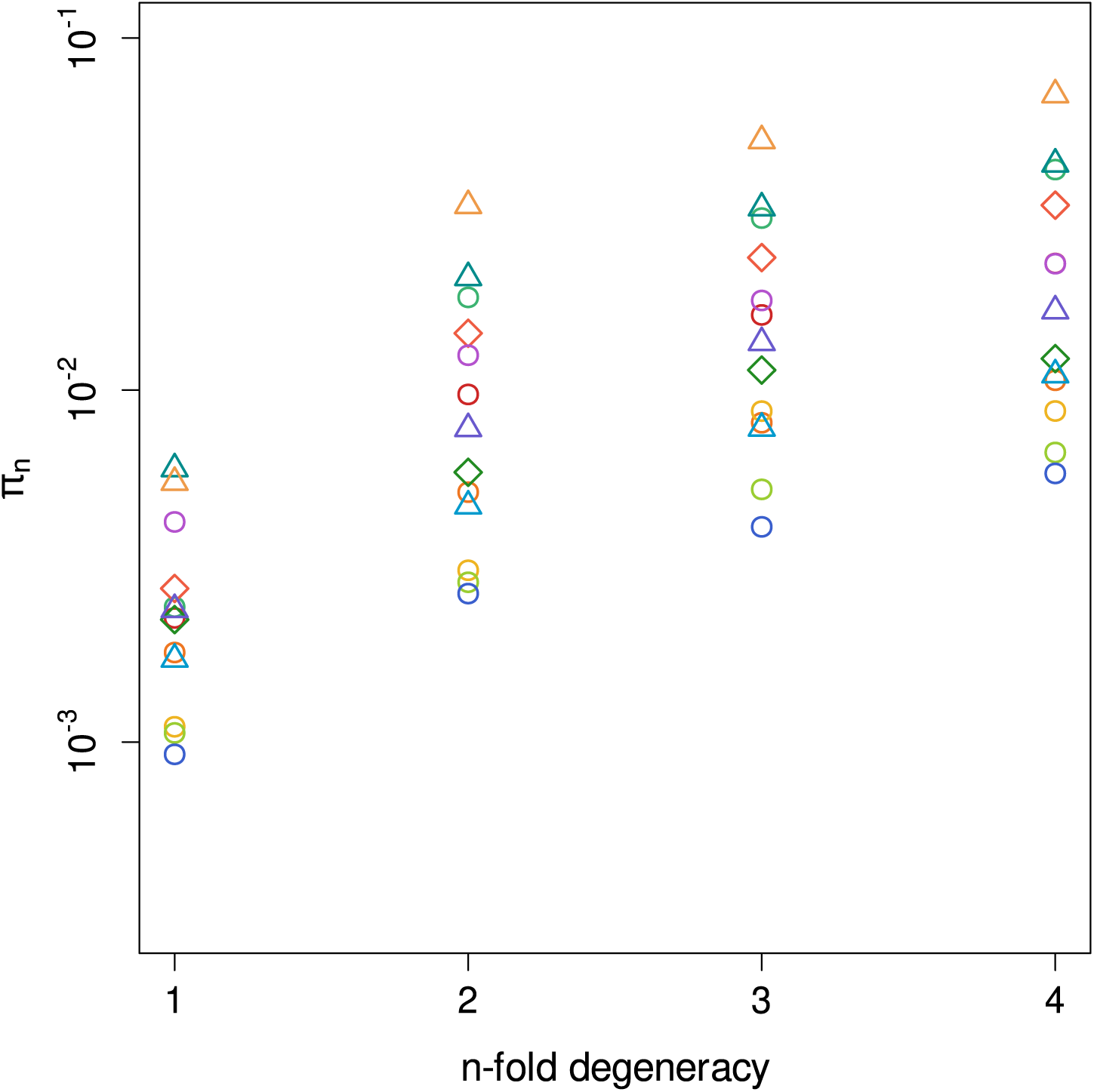
Increase in nucleotide diversity with n-fold degeneracy of codon sites across 13 species. Species are shown in different colors and symbols as defined in the legend of main Figure 2.

**Figure S5:**
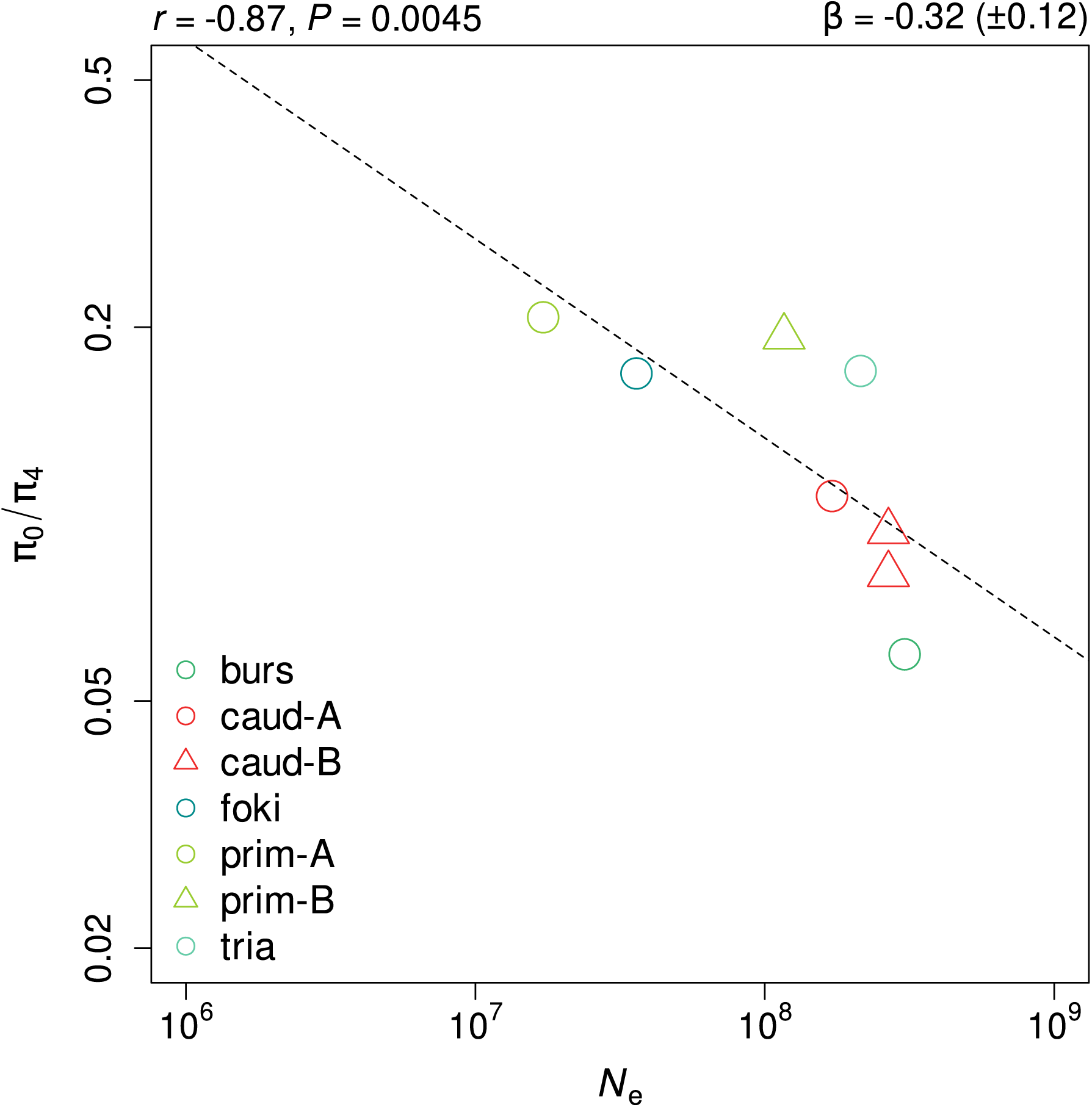
Correlation between *π*_0_*/π*_4_ and *N_e_* estimated using separate populations, for species with at least 2 populations. Dashed line shows the linear regression fit on log-log scale, with slope (*β*) and standard error given on top-left, and Spearman correlation (*r*) coefficient on the top-right.

**Figure S6:**
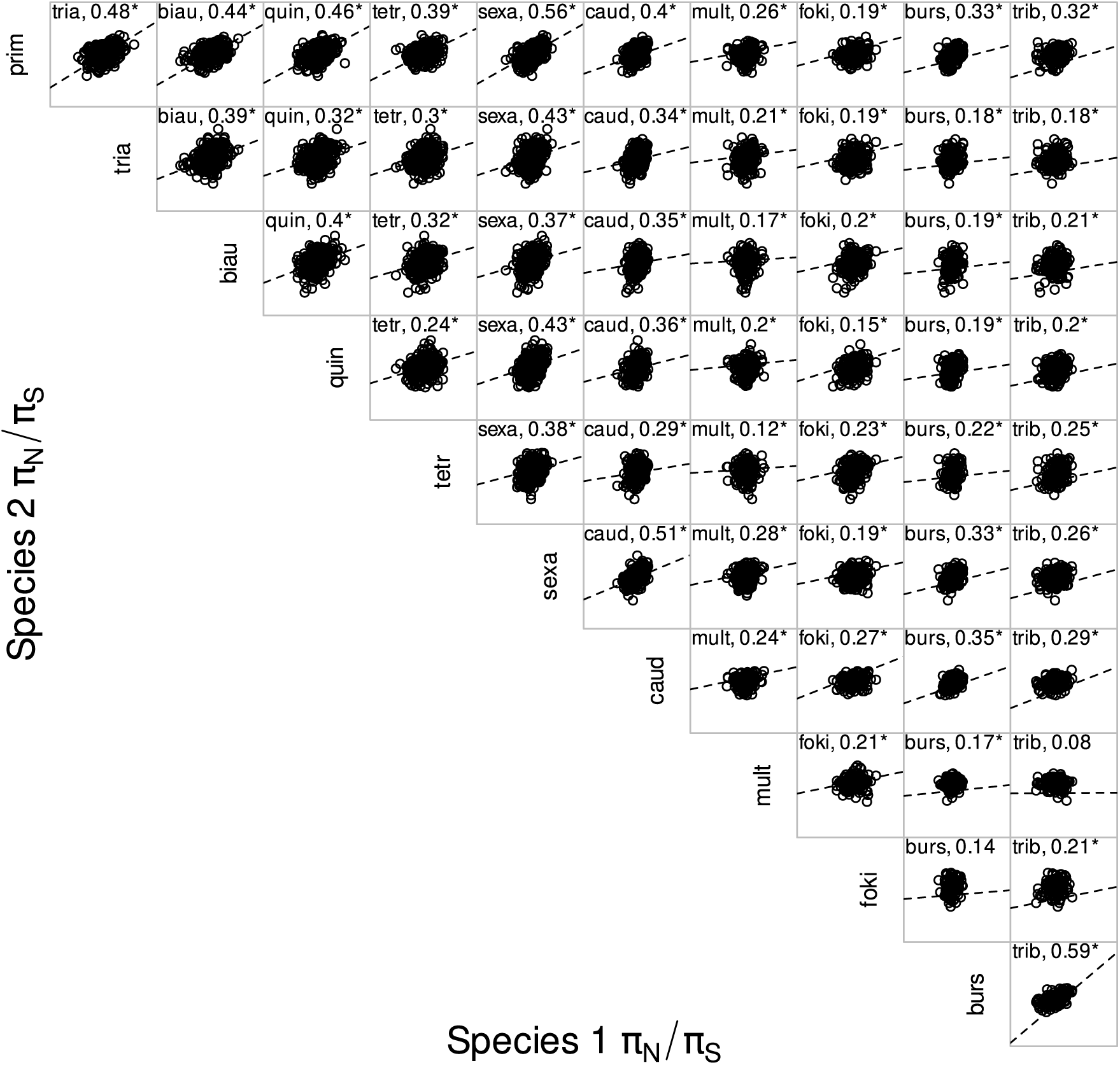
Scatterplots of pairwise comparisons in *π_N_ /π_S_* among 11 species of *Paramecium*, for 717 core macronuclear proteins of *P. aurelia*. The corresponding Spearman correlation coefficient is shown in each panel, with those marked with an asterisk having *P <* 0.05. Dashed lines show linear regression fit on a log-log scale.

